# Age-dependent H3K9 trimethylation by dSetdb1 impairs mitochondrial UPR leading to degeneration of olfactory neurons and loss of olfactory function in *Drosophila*

**DOI:** 10.1101/2024.06.17.599276

**Authors:** Francisco Muñoz-Carvajal, Nicole Sanhueza, Mario Sanhueza, Felipe A. Court

**Affiliations:** Center for Integrative Biology, Faculty of Sciences, Universidad Mayor, Santiago, Chile; Geroscience Center for Brain Health and Metabolism (GERO), Santiago, Chile; Center for Resilience, Adaptation and Mitigation, Universidad Mayor, Temuco, Chile; Buck Institute for Research on Aging, Novato, CA, USA

**Keywords:** mitochondrial unfolded protein response, H3K9me3, neurodegeneration, aging, mitochondria, methylation, olfaction

## Abstract

Aging is characterized by a decline in essential sensory functions, including olfaction, which is crucial for environmental interaction and survival. This decline is often paralleled by the cellular accumulation of dysfunctional mitochondria, particularly detrimental in post-mitotic cells such as neurons. Mitochondrial stress triggers the mitochondrial unfolded protein response (UPR^MT^), a pathway that activates mitochondrial chaperones and antioxidant enzymes. Critical to the efficacy of the UPR^MT^ is the cellular chromatin state, influenced by the methylation of lysine 9 on histone 3 (H3K9). While it has been observed that the UPR^MT^ response can diminish with an increase in H3K9 methylation, its direct impact on age-related neurodegenerative processes, especially in the context of olfactory function, has not been clearly established. Using *Drosophila,* we demonstrate that an age-dependent increase in H3K9 trimethylation by the methyltransferase dSetdb1 reduces the activation capacity of the UPR^MT^ in olfactory projection neurons leading to neurodegeneration and loss of olfactory function. Age-related neuronal degeneration was associated with morphological alterations in mitochondria and an increase in reactive oxygen species levels. Importantly, forced demethylation of H3K9 through knockdown of dSetdb1 in olfactory projection neurons restored the UPR^MT^ activation capacity in aged flies, and suppressed age-related mitochondrial morphological abnormalities. This in turn prevented age-associated neuronal degeneration and rescued age-dependent loss of olfactory function. Our findings highlight the effect of age-related epigenetic changes on the response capacity of the UPR^MT^, impacting neuronal integrity and function. Moreover, they suggest a potential therapeutic role for UPR^MT^ regulators in age-related neurodegeneration and loss of olfactory function.

## Introduction

Aging is associated to a time-dependent organ dysfunction that increases the vulnerability of an organism to various forms of stress, ultimately leading to organism death^1–7^. Aging induces alterations across various physiological systems, with the olfactory system being particularly affected^1^. Olfaction, the sense of smell, is essential for detecting environmental odors crucial for feeding, reproductive and survival behaviors^2,3^. Importantly, dysfunction in olfaction is emerging as one of the early signs of neurodegenerative diseases, including Alzheimer’s and Parkinson’s disease^8,9,11^. Research on *Drosophila melanogaster* has shown that the age-related decline in odor response is influenced by the functional state of mitochondria in olfactory projection neurons (OPNs)^4^. This decline is accompanied by defects in neuronal integrity and a decrease in synaptic proteins^5^, which correlates with a reduction in mitochondria and an increase in ROS^5^, but the underlying mechanisms has not been defined.

Under compromised mitochondrial integrity or function, cells engage in a transcriptional response known as the mitochondrial unfolded protein response (UPR^MT^)^14^. This mitochondrial program can be activated by the impairment of the electron transport chain (ETC), alteration of mitochondrial dynamics, accumulation of unfolded proteins, reduction of mitochondrial DNA, or reduction of mitochondrial chaperones or protease^6–9^,. Upon UPR^MT^ pathway engagement, the transcription factor ATFS-1 (*C. elegans* ortholog of mammalian ATF5 and *Drosophila* crc) translocates from the mitochondria to the nucleus^10,11^. In the nucleus, ATFS-1, DVE-1, and UBL-5 interact to reorganize the chromatin structure, enabling activation of the nuclear transcription of mitochondrial chaperones, including hsp-60 and hsp-6, and the protease clpp-1. This coordinated transcriptional response restores mitochondrial function under stress conditions by metabolic adaptations, and enhancing mitochondrial biogenesis^12–14^.

Chromatin remodeling is crucial for UPR^MT^ regulation, with the epigenetic state of lysine 9 on histone 3 (H3K9) serving as a critical regulator of the response^14–17^. Changes in H3K9 methylation by the methyltransferase MET-2, the *C. elegans* ortholog of human SETDB1, modify UPR^MT^-related loci exposure, modulating binding of UPR^MT^ regulators DVE-1 and ATFS-1^21^. In addition, enzymes that remove methyl groups from H3K9 significantly influence UPR^MT^ activation. For example, demethylases JMJD-3.1 and JMJD-1.2 remove trimethylation from H3K9me3 and H3K27me3, enabling UPR^MT^ activation^15,17^. Recent studies revealed the effects of H3K9me3 methylation on UPR^MT^ activation and mitochondrial function across species. In C. *elegans* the epigenetic factors BAZ-2 and SET-6, which regulate H3K9me3 levels, have conserved roles in impacting aging processes through mitochondrial function^18^. In mice, deletion of Baz2b, a homologue of BAZ-2, shows beneficial effects on mitochondrial function and cognitive abilities, indicating a conserved mechanism across species that influences aging and healthspan through mitochondrial function and epigenetic regulation^18^. Accordingly, in the hippocampus of aged mice, H3K9me3 levels rise with age, a change associated with age-related cognitive decline^19^. Age-related increases in H3K9me3 have also been observed in the brains of aged *Drosophila* and muscle stem cells of aged mice^20,21^. While previous research has linked changes in methylation levels to age-related functional decline, the specific role of these epigenetic alterations in the context of neuronal degeneration through reduced UPR^MT^ activation capacity remains to be fully elucidated. This study aims to bridge this gap by providing detailed insights into how epigenetic mechanisms, particularly methylation changes, directly contribute to the aging-associated loss of olfactory function and neuronal degeneration by impacting the UPR^MT^ pathway and mitochondrial function.

Here, we employed behavioral, molecular, and morphological methodologies to investigate whether epigenetic regulation of UPR^MT^ is linked to neurodegeneration in the aging brain and its involvement in age-associated olfactory decline. To this end, we utilized the OPNs in the adult *Drosophila* antennal lobe (AL), which exhibit age-related neurodegeneration correlating with functional neuronal decline^5^. Our results demonstrate that with aging, there is a decline in the response capacity of the UPR^MT^ in OPNs, functionally associated with a dSetdb1-dependent increase in H3K9me3 levels. Genetic inhibition of dSetdb1 reduces H3K9me3 levels, enabling the activation of UPR^MT^, restoring mitochondrial oxidation to youthful levels, and preventing age-associated degeneration of OPNs. This effect is particularly evident in the somas located in the antennal lobe (AL) and the presynaptic connections of the lateral horn (LH) in the *Drosophila* brain. Importantly, maintaining UPR^MT^ activation during aging preserved olfactory function. These findings underscore the critical role of epigenetic regulation, specifically through dSetdb1 and H3K9me3, in modulating neuronal integrity and sensory function during aging.

## Results

### 1. UPR^MT^ response capacity decreases with aging in the *Drosophila* antennal lobe

To study the modulation of UPR^MT^ along aging and its association to olfactory function, we first generated reporters based on the expression of the fluorescent protein dsRed under the promoters of chaperones hsp60, which specifically responds to UPR^MT^ stimuli in across species and has been effectively used to indicate UPR^MT^ activation^22–26^. We focused on the antennal lobe (AL), the functional homolog of the vertebrate olfactory bulb, where olfactory projection neurons process olfactory sensory inputs. In young flies (0 dpe), a low-intensity signal from the Hsp60::dsRed reporter was detected under control conditions, which significantly increased upon exposure to the UPR^MT^ activator paraquat (PQ, Fig. 1A and B). Importantly, this increase in Hsp60::dsRed signal was not induced by non-specific mitochondrial stressor tunicamycin, which activates UPR^ER^ and the ER-stress specific reporter Xbp1::GFP^27^ (Fig. 1A-D). To further evaluate the specificity of the UPR^MT^ reporter, we downregulated the UPR^MT^ nuclear activators *dve*, *ubl,* and *crc*. Pan-neuronal downregulation of these UPR^MT^ activators significatively reduced the Hsp60::dsRed response to PQ compared to control flies (Fig. 1E and F), indicating that this novel reporter can be used to specifically monitor UPR^MT^ activity. We next used the Hsp60::dsRed reporter to evaluate UPR^MT^ activity during aging in the *Drosophila* AL. Compared to the robust signal triggered by PQ in young flies, aged flies (45 dpe) did not exhibit Hsp60::dsRed reporter activation in AL neurons after PQ (Fig. 1G-H). Importantly, no significant changes in GFP-labelled neuronal volume were observed in aged versus young flies or after PQ treatment (Fig. 1I). This data suggests that the ability to trigger UPR^MT^ activity declines with advanced age.

**Figure 1.**
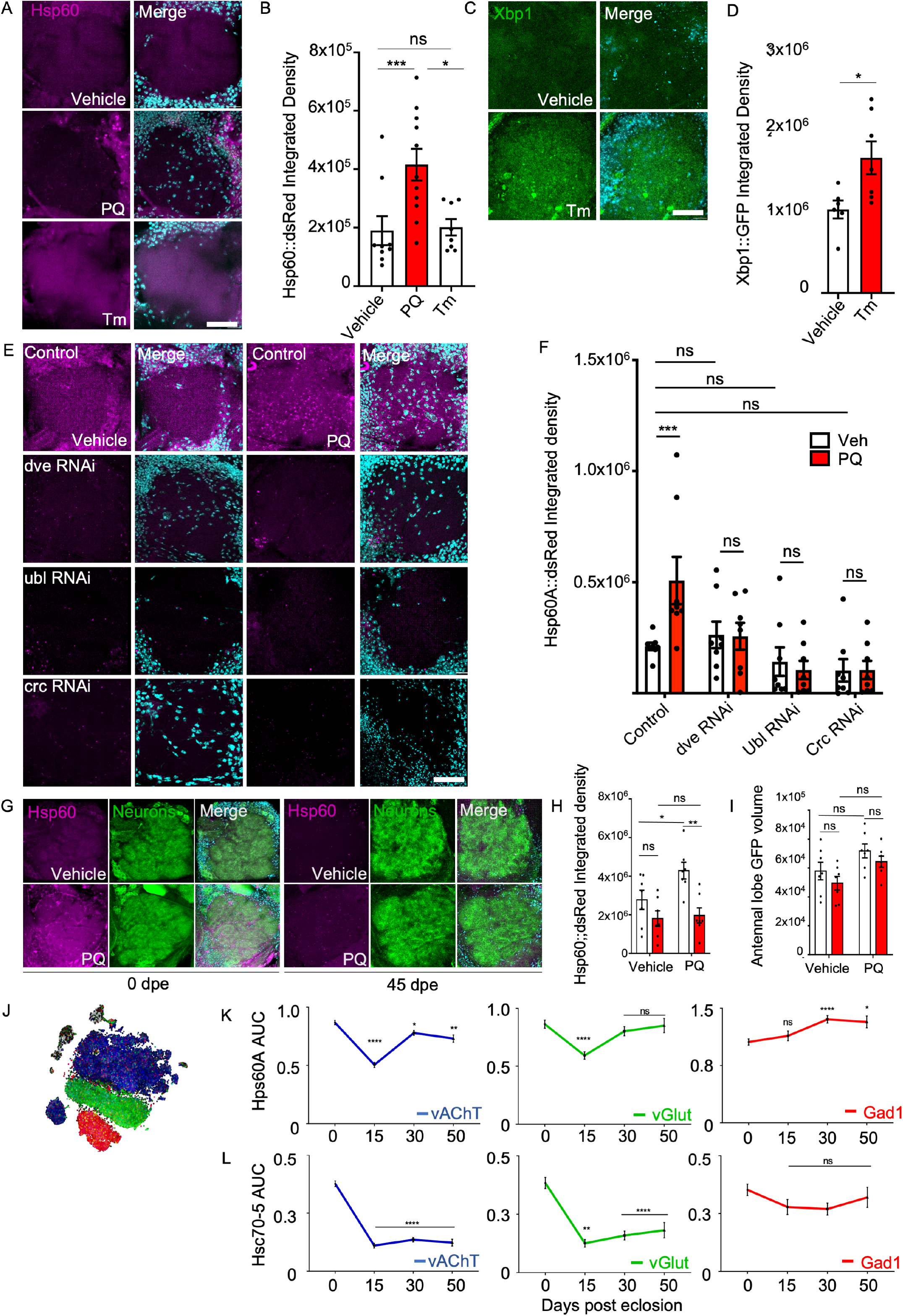
UPR^MT^-dependent activation of the Hsp60::dsRed reporter in the Antennal Lobe (AL) of *Drosophila*. (A) Representative images of AL from UPR^MT^ reporter flies at 0 dpe treated with either 10 mM Paraquat (PQ), 0.25 µg/µl Tunicamycin (Tm) for 48 hours, or vehicle (dH2O). Magenta indicates the Hsp60::dsRed signal, with cyan representing the DAPI nuclear signal. Scale bar, 20 µm. (B) Bar plot of integrated density of the Hsp60::dsRed signal for the conditions shown in panel A. Two-way ANOVA Bonferroni’s test: vehicle vs. PQ (p=0.0061, n1=9, n2=11), PQ vs. Tm (p=0.0119, n1=11, n2=8), vehicle vs. Tm (p=0.9863). n=Antennal lobe of Drosophila Brain. (C) Representative images of AL from Xbp1::GFP flies treated with 0.25 µg/µl Tm for 48 hours. The Xbp1::GFP signal appears in green, with cyan for DAPI. Scale bar, 20 µm. (D) Integrated density measurement of the Xbp1::GFP signal for the conditions in panel C. Unpaired t-test: Tm vs. vehicle (p=0.0231, n1=7, n2=8).n=Antennal lobe of *Drosophila* Brain. (E) Hsp60::dsRed in AL with pan-neuronal RNAi knockdown of dve, ubl, crc, post treatment with PQ or vehicle, magenta shows the Hsp60A-dsRed signal, cyan for DAPI. Scale bar, 20 µm. (F) Integrated density of Hsp60::dsRed, Two-way ANOVA Bonferroni’s tes between veh vs PQ treated flies: Control (p=0.0038, n=8), dve (p>0.9999, n=8), crc (p>0.9999, n=8), ubl (p>0.9999, n=8),. (G) Representative images Hsp60::dsRed reporter in GFP labeled neurons of the AL of flies treated with 10mM PQ or vehicle for 48 hours. Hsp60::dsRed in magenta, Pan-neuronal GFP in green, and DAPI in cyan. Scale bar, 20 µm. (H) Neuronal volume (µm ^3^) in AL from 0 to 45 dpe. Two-way ANOVA with Dunnett’s multiple comparisons between 0 and 45 dpe flies: for vehicle (p=0.4255, n=7) and PQ (p=0.5076, n=7). Comparing vehicle to PQ treatment showed no significant difference at 0 dpe (p=0.1133, n=7) and 45 dpe (p=0.0857, n=7). (I) Integrated density of Hsp60::dsRed in GFP-labeled neurons from 0 to 45 dpe. Two-way ANOVA with Dunnett’s multiple comparisons between ages for vehicle (p=0.2319, n=7) and for PQ treatment (p=0.0017, n=7). For difference between treatments: vehicle and PQ at 0 dpe (p=0.0404, n=7), and at 45 dpe (p=0.9581, n=7). n=Antennal lobe of Drosophila Brain (J) Dot plot visualizing vAChT (blue), vGlut (green), and Gad1 (red) for UPR^MT^ activation analysis via single-cell RNA seq data. (E) AUC scores of Hsp60 expression: vAChT (0 vs. 50 dpe, p=0.0049, n=576). vGlut (0 vs. 50 dpe, p=0.9998, n=168). Gad1 (0 vs. 50 dpe, p=0.0191, n=168). (F) AUC scores of Hsc70-5 expression: vAChT (0 vs. 50 dpe, p<0.0001, n=576). vGlut (0 vs. 50 dpe, p<0.0001, n=168). Gad1 (0 50 dpe, p=0.9914, n=168). n= Singel cell expression. Statistical significance is denoted as **** p < 0.0001; *** p < 0.001; ** p < 0.01; * p<0.05; ns > 0.05.

We then explore the age-dependent endogenous expression of UPR^MT^-associated chaperones Hsp60 and Hsc70-5 using Scope, a single-cell gene expression repository of brain cells from *Drosophila* at different ages (http://scope.aertslab.org)^28^. The *Drosophila* brain consists of three major groups of neurons: glutamatergic, GABAergic and cholinergic neurons, with the latter being the most abundant in the AL (Fig. 1J). Single-cell data for cholinergic neurons show that both Hsp60A and Hsc70-5 expression levels decrease in aged flies compared to young animals (Fig. 1K-L). Interestingly, this was also observed in glutamatergic neurons but not in GABAergic neurons, suggesting differences in UPR^MT^ activation in different neuronal populations. These findings suggest that in the *Drosophila* AL, aging is associated with a decline in UPR^MT^ activity and chaperone expression, particularly in cholinergic neurons.

### 2. Epigenetic regulation of the UPR^MT^ by dSetdb1 in the AL of *Drosophila* brain

It has been previously demonstrated that trimethylation of H3K9 increases during *Drosophila* aging^20^, a phenomenon that mirrors observations in other species. To assess whether methylation levels of H3K9 can modulate UPR^MT^ activation in flies, we studied flies with pan-neuronal knockdown of dSetdb1, a specific H3K9 methyltransferase. Our data demonstrates that pan-neuronally downregulating dSetdb1 prevents the age-associated increase in H3K9 trimethylation in homogenates of *Drosophila* heads (Fig. 2A). We then investigated the role of H3K9 trimethylation in UPR^MT^ activation by examining the Hsp60::dsRed reporter in flies with a ubiquitous loss of function of dSetdb1. Control flies exhibited similar basal levels of Hsp60::dsRed signal in the AL of young and aged flies. Similarly, we observed no differences in the reporter signal for young flies in which dSetdb1 was downregulated (Fig. 2B), consistent with the low levels of H3K9 trimethylation observed in young animals (Fig. 2A). However, the Hsp60::dsRed signal in aged dSetdb1 mutants was significantly higher compared to age-matched control flies (Fig. 2B-C). These results were further confirmed using a second UPR^MT^ reporter based on Hsc70-5 expression (Fig. 2D-E). These data suggest that dSetdb1 contributes to age-dependent H3K9 trimethylation, and its reduced function in the AL of aged flies correlates with a basal increase in UPR^MT^ activity.

**Figure 2.**
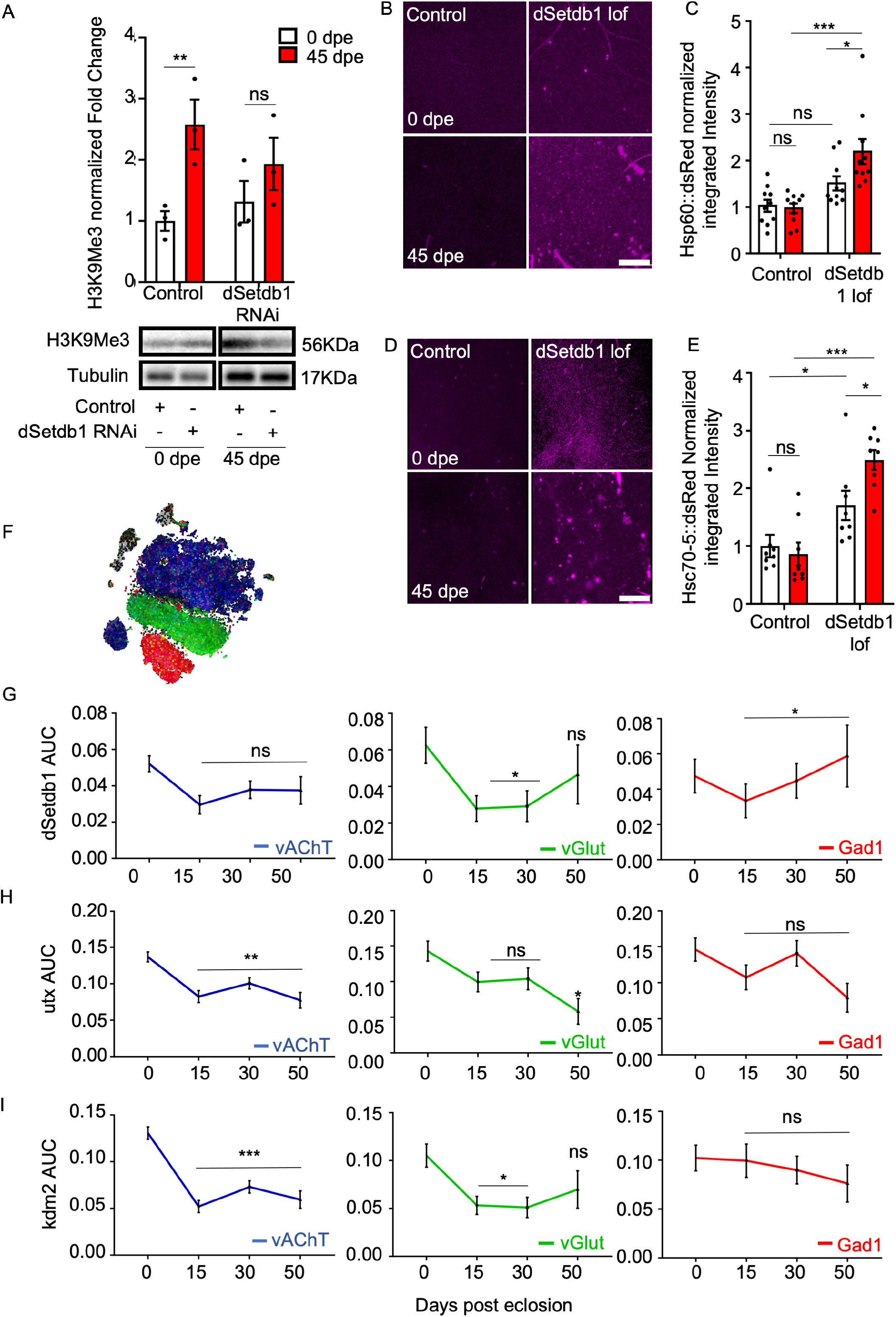
dSetdb1 negatively regulates UPR^MT^ in aging through increasing H3K9me3 levels in AL of *Drosophila*. (A) Western blot for H3K9me3 levels with pan-neuronal downregulation of dSetdb1. Two-way ANOVA Bonferroni’s multiple comparisons test results: Control at 0 vs. 45 dpe (p=0.0255, n=3), dSetdb1 RNAi at 0 vs. 45 dpe (p=0.4951, n=3), Control vs. dSetdb1 RNAi at 0 dpe (p>0.9999, n=3), and at 45 dpe (p=0.4556, n=3). n = 20 fly heads. (B) Representative confocal images of AL from UPR^MT^ reporter flies with dSetdb1 loss of function, displaying Hsp60A::dsRed in magenta. Scale bar, 20 µm. (C) Normalized integrated density of Hsp60A::dsRed. Two-way ANOVA Bonferroni’s multiple comparisons test between 0t vs. 45 dpe within dSetb1 Lof genotype (p<0.0001, n=10) Control (p >0.9999, n=10). (D) Confocal images of Hsc70-5::dsRed in AL from flies at 0 and 45 dpe with dSetdb1 loss of function. Scale bar, 20 µm. (E) Integrated density of Hsc70-5::dsRed normalized to control values. Two-way ANOVA Bonferroni’s multiple comparisons between 0 vs. 45 dpe within dSetdb1 Lof genotype (p<0.0001, n=8) and control (p=0.8633, n=8). n= AL of Drosophila Brain. (F) Dot plot from Scope single-cell RNA-seq analysis depicting neuronal types: vAChT (blue), vGlut (green), and Gad1 (red). (G) AUCell scores for dSetdb1 expression in single neurons at 0, 15, 30 and 50dpe. One-way ANOVA with Dunnett’s multiple comparison against 0 dpe. vAChT neurons: 50 dpe (p=0.2906, n1=2932, n2=656). vGlut neurons: 50 dpe (p=0.7386, n1=712, n2=172). Gad1 neurons: 50 dpe (p=0.8916, n1=576, n2=197). (H) AUCell scores for utx expression in single neurons at 0, 15, 30 and 50dpe. One-way ANOVA with Dunnett’s multiple comparison against 0 dpe. vAChT neurons: 50 dpe (p=0.0001, n1=2932, n2=656). vGlut neurons: 50 dpe (p=0.321, n1=712, n2=172). Gad1 neurons: 50 dpe (p=0.0688, n1=576, n2=197). (I) AUCell scores for kdm2 expression in single neurons at 0, 15, 30 and 50dpe. One-way ANOVA with Dunnett’s multiple comparison against 0 dpe. vAChT neurons: 50 dpe (p<0.0001, n1=2932, n2=656). For vGlut neurons: 50 dpe (p=0.321, n1=712, n2=172). For Gad1 neurons: 50 dpe (p=0.667, n1=576, n2=197). All error bars represent SEM. P-value: **** p < 0.0001; *** p < 0.001; ** p < 0.01, * p<0.05 and ns > 0.05.

To understand the relevance of H3K9me3-related genes in a neuron-specific context, we then analyzed single-cell data from Scope to assess the expression levels of dSetdb1, as well as the H3K9 demethylases Utx and Kdm2^15,29^. In vAChT neurons, dSetdb1 expression remains constant throughout aging (Fig. 2G). However, both Utx and Kdm2 exhibit an age-dependent decrease in expression (Fig. 2H and I). Together, this data suggests that the age-dependent reduction in H3K9 demethylation enzymes could be associated with higher levels of H3K9me3 in the aged *Drosophila* brain, which in turn might contribute to the age-related decrease in UPR^MT^ activity.

### 3. Age-dependent decline in olfactory function depends on the epigenetic modulation of the UPR^MT^

Having established that the decline in UPR^MT^ activity in aged *Drosophila* is linked to elevated levels of H3K9me3, we then explored the potential link between UPR^MT^ activation and olfactory function in *Drosophila*. The ability to discern between odors diminishes with age in flies, a phenomenon quantifiable through the olfactory T-maze (Fig. 3A-B). Therefore, we genetically downregulated the UPR^MT^ transcriptional activators dve, ubl or crc and studied the ability of flies to discriminate odors throughout their lifespan. Remarkably, young flies with the knockdown of the nuclear activators of the UPR^MT^ exhibited a reduced olfactory capacity to discriminate an abrasive odor compared to control animals (Fig. 3C). Aged flies with the knockdown of dve, ubl or crc did not show significantly different olfactory function compared to age-matched controls or to young flies from the same genotype (Fig. 3C). These findings underscore the crucial role of UPR^MT^ transcriptional activators in preserving olfactory discrimination. We next explored the impact of the epigenetic regulation of UPR^MT^ in neuronal functionality. To this end, we generated flies with pan-neuronal knockdowns of the H3K9 methyltransferase dSetdb1 and demethylases Kdm2 or Utx. Consistent with our previous observations, young flies with downregulated dSetdb1 did not show a difference in H3K9me3 levels compared to controls. In contrast, downregulation of demethylases Utx or Kdm2 led to increased H3K9me3 levels, highlighting their distinct regulatory roles (Fig. 3D). To determine if H3K9me3 levels alter neuronal functionality in the olfactory system, we assessed olfactory function in flies with downregulation of dSetdb1, Utx, or Kdm2. In young flies, pan-neuronal knockdown of dSetdb1 showed no significant difference from control flies. However, aged dSetbd1 mutant flies exhibited improved olfactory function, with no significant difference when compared with young flies (Fig. 3E). On the other hand, pan-neuronal knockdown of Utx or Kdm2 impaired olfactory function in young flies, with an odor discrimination capacity similar to aged control flies (Fig. 3E), indicating that age-dependent increases in H3K9me3 progressively affects olfactory function.

**Figure 3.**
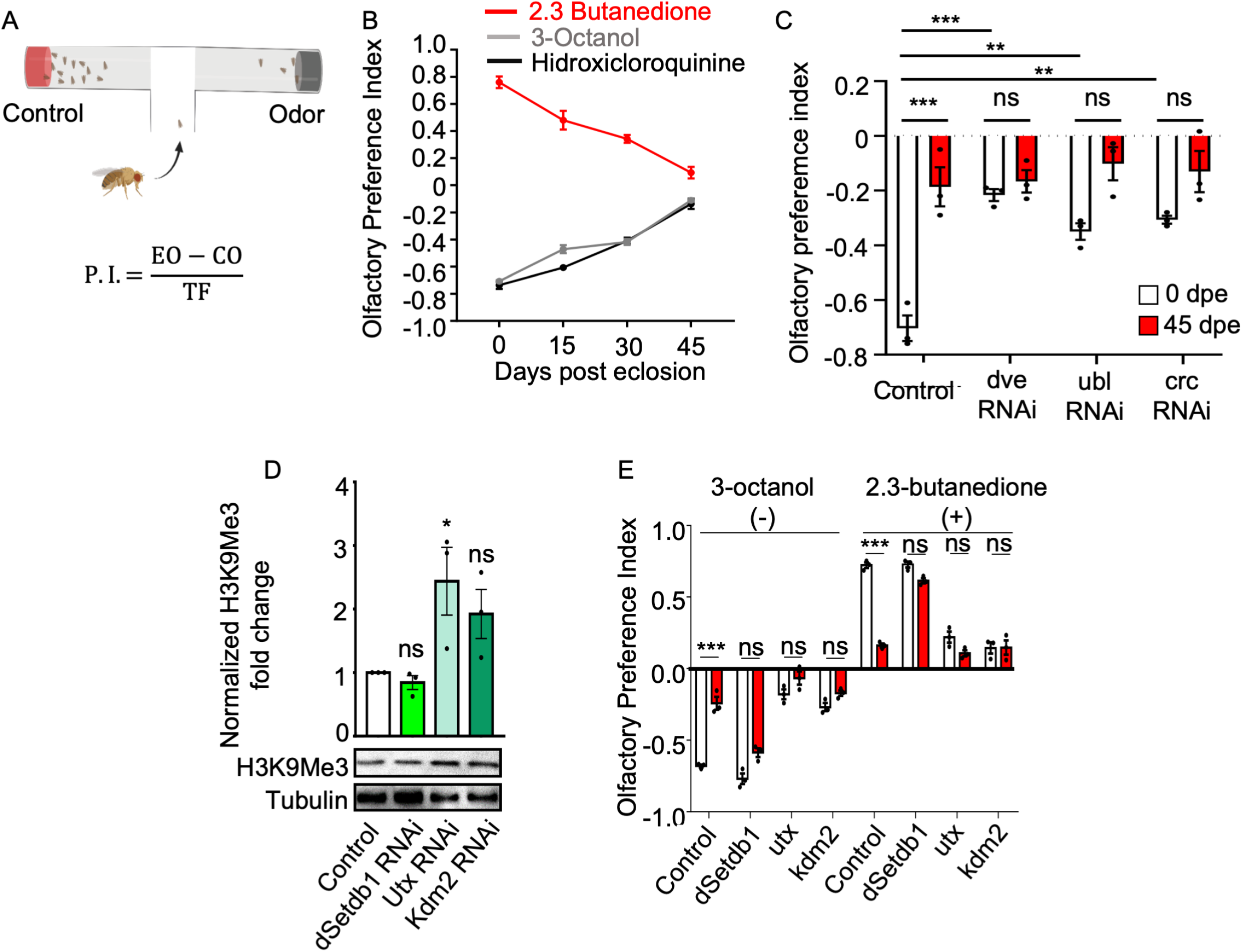
dSetdb1 pan-neuronal downregulation preserves olfactory function in aging. (A) Olfactory T-maze was used to perform the olfactory behavioral test. Flies are presented to an experimental odor or vehicle. Flies have 60 seconds to discriminate between odors and go to an arm of the T-maze. At the end of the time, an image is acquired, and the preference index is calculated for every trial; every dot corresponds to 10 trials of 15 flies each. (B) Olfactory preference index shows the aging-associated functional decline in the olfactory system. (C) Olfactory preference index in flies with pan-neuronal downregulation of dve, ubl, and crc. n = 3 populations of 10 flies each. (D) Western blot analysis of H3K9me3 levels in young flies with pan-neuronal downregulation of dSetdb1 (green), utx (gray), and kdm2 (calypso). Each n represents homogenized pools of 20 fly heads. One-way ANOVA Bonferroni’s multiple comparisons test results: Control vs. dSetdb1 RNAi (p=0.9738, n=3), Control vs. utx RNAi (p=0.0391, n=3), Control vs. kdm2 RNAi (p=0.1951, n=3). (E) Olfactory preference indices for 0 and 45 days post-eclosion (dpe) flies with downregulation of dSetdb1, utx, and kdm2, exposed to odors 3-octanol (-) and 2,3-butanedione (+), revealed the following Two-way ANOVA Bonferroni’s test results: For 3-octanol (-), control at 0 vs. 45 dpe (p<0.0001, n=3), dSetdb1 (p=0.0624, n=3), utx (p=0.1356, n=3), kdm2 (p=0.298, n=3); for 2,3-butanedione (+), control (p<0.0001, n=3), dSetdb1 (p=0.1356, n=3), utx (p=0.1374, n=3), kdm2 (p>0.9999, n=3). Each n represents populations of 10 flies.

### 4. Epigenetic modulation of the UPR^MT^ influences olfactory function in an OPN-cell autonomous manner

As olfactory function is a complex behavior dependent on multiple central and peripheral neuronal populations, we investigated whether age-related changes in H3K9 methylation, UPR^MT^ and olfactory function were specifically associated with cholinergic OPNs. We first assessed age-dependent changes in H3K9me3 trimethylation in olfactory projection neurons (OPNs). To this end, we selectively label the plasma membrane of OPNs by expressing the CD8::GFP fusion protein using the OPN-specific driver GH146-Gal4. We then performed immunofluorescence to evaluate H3K9me3 levels specifically in ToPro3-positive nuclei located in GFP-positive neurons. Using this method, we observed an increase in trimethylation in aged OPNs compared to young ones (Fig. 4A-C). We then evaluated the regulation of dSetdb1 in aging-associated OPNs trimethylation by generating flies carrying the knockdown of dSetdb1 specifically in cholinergic OPNs. Remarkably, H3K9 trimethylation levels in aged flies with OPN-specific dSetdb1 knockdown were reversed and with no significant difference from young control flies (Fig. 4A-C).

**Figure 4.**
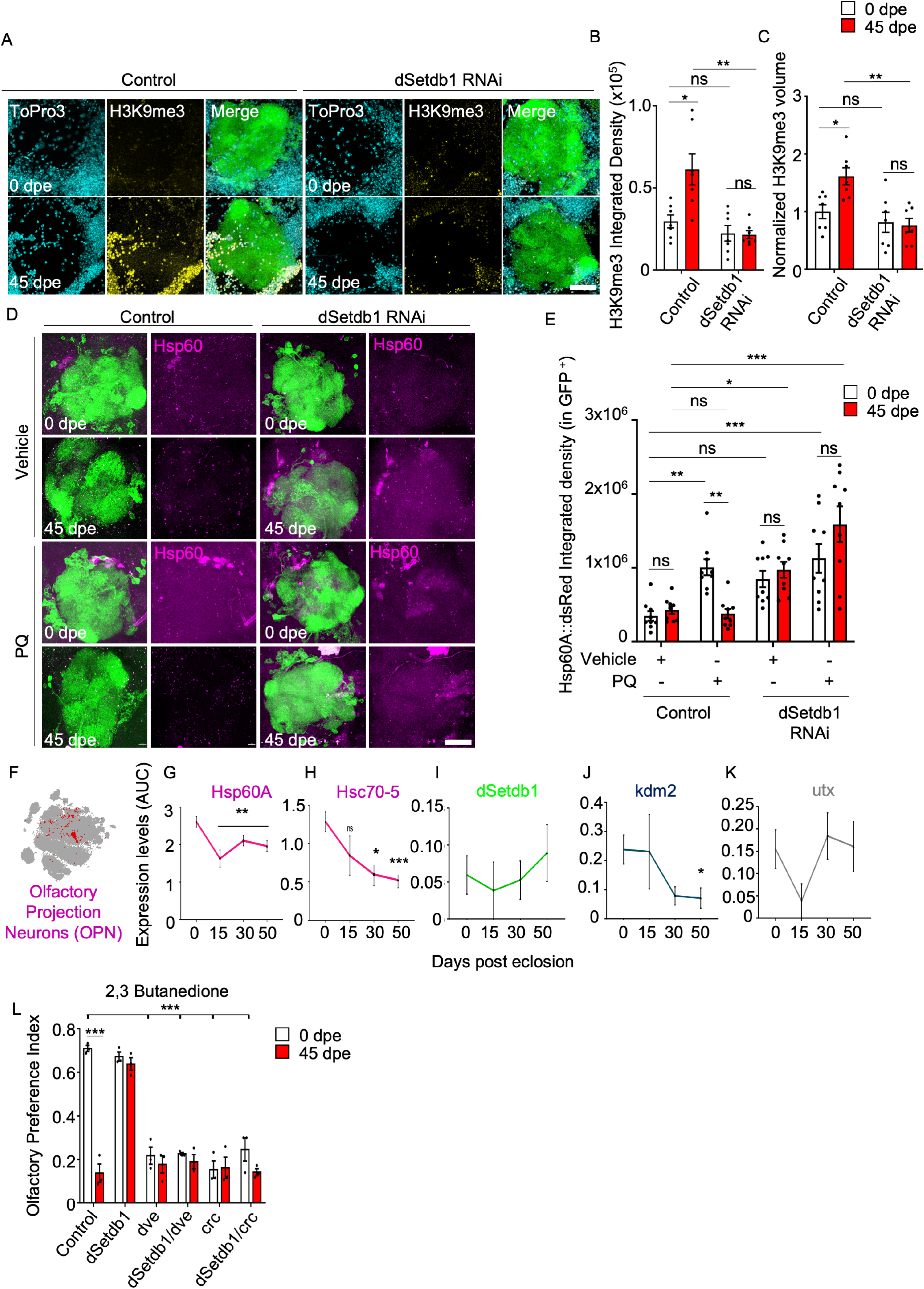
dSetdb1 Downregulation Preserves OPNs Function in Aging by Reducing H3K9me3 and enabling UPR^MT^. (A) Representative images of GFP-tagged OPNs in the AL of 0 and 45 dpe flies bearing the downregulation of dSetdb1 under the control of the GH146 driver. Nuclei are in cyan (ToPro3), H3K9me3 is in yellow, and the right panel is a merge of three channels with OPNs in green. Scale bar, 20 µm. (B) Analysis of H3K9me3 integrated density in the nuclei of GFP-tagged OPNs across aging in *Drosophila* with dSetdb1 knockdown. Using two-way ANOVA with Bonferroni’s correction, control flies demonstrated a significant reduction in density from 0 to 45 dpe (p=0.0128, n=7), while dSetdb1 RNAi flies (p=0.5022, n=7), between control vs dSetdb1 RNAi at 0 dpe (p=0.3413, n=7). At 45 dpe, dSetdb1 RNAi vs control (p=0.0088, n=7). (C) Quantification of H3K9me3 volume, specifically in nuclei of GFP signal, normalized to control of 0 dpe untreated flies. The white bars represent 0 dpe flies and red bars 45 dpe flies. n=7. Two-way ANOVA Bonferroni’s multiple comparisons showed: For control flies of 0 vs 45 dpe p=0.0057. For dSetdb1 RNAi group of 0 vs 45dpe (p>0.9999), Control vs dSetdb1 RNAi at 0 dpe p=0.0604, for 45 dpe, Control vs dSetdb1 RNAi p=0.0004, n= independent AL of *Drosophila* brain. (D) Representative images of OPNs labeled with GFP (green) bearing the UPR^MT^ reporter Hsp60::dsRed (magenta) in the AL of 0 and 45dpe flies treated with PQ or vehicle for 48h. Scale Bar, 20µm. (E) Quantification of Integrated density of Hsp60A::dsRed specifically in the GFP labeled neurons in the AL of 0 and 45 dpe flies treated with vehicle or PQ 10mM for 48h. Two-way ANOVA Bonferroni’s test result at 0 dpe between Control Veh vs. Control PQ p= 0.0027, n=9; Control Veh vs. dSetdb1 Veh p= 0.0291, n=9; Control Veh vs. dSetdb1 PQ p= 0.0003, n=9 and at 45 dpe between Control Veh vs. Control PQ p= 0.9872, n=9; Control Veh vs. dSetdb1 Veh p= 0.0154, n=9; Control Veh vs. dSetdb1 PQ p<0.0001, n=9. n= AL of *Drosophila* brain. (F) Dot plot of expression cluster showing specifically OPN cluster in red. One-way ANOVA with Dunnett’s multiple comparisons test against 0 dpe was performed for (G) Hsp60A Expression: At 0 vs 50 dpe p=0.0102, n1=60, n2=38.and (I) Hsc70-5 expression 0 vs 50 dpe p<0.0001, n1=60, n2=38. (I-K) Expression levels of dSetdb1, kdm2, and utx in OPNs through aging. One-way ANOVA with Dunnett’s multiple comparisons test against 0 dpe was performed., (I): dSetdb1 Expression 0 vs 50 dpe (p=0.0102, n1=84, n2=56). (J) utx Expression 0 vs 50 dpe (p=0.9996, n1=84, n2=56). (K) kdm2 0 vs 50 dpe (p=0.0425, n1=84, n2=56). Each n represents the AUC values for expression in a single cell. (L) Olfactory preference index of flies bearing the GH146 Gal4 driven knockdown of dSetdb1, dve. crc and the double knockdown of dSetdb1/dve and dSetdb1/crc, respectively. White bars are 0 dpe flies and red bars are 45 dpe flies. n = 3 population of 10 flies. Results from a Two-way ANOVA Bonferroni’s multiple comparisons test are as follows: Control at 0 vs. 45 dpe: p<0.0001, n=3, dSetdb1 at 0 vs. 45 dpe: p>0.9999, n=3, dve at 0 vs. 45 dpe: p>0.9999, n=3, dSetdb1/dve at 0 vs. 45 dpe: p>0.9999, n=3, crc at 0 vs. 45 dpe: p>0.9999, n=3, dsetdb1/crc at 0 vs. 45 dpe: p=0.2606, n=3. P-value: **** p < 0.0001; *** p < 0.001; ** p < 0.01, * p<0.05 and ns > 0.05.

To further explore the neuronal-specificity of the UPR^MT^ effect, we assessed Hsp60::dsRed reporter activity specifically in aged OPNs. The response of the UPR^MT^ sensor in CD8::GFP tagged neurons increased in young flies treated with PQ compared to animals treated with vehicle. However, this response to the mitochondrial stressor was diminished in aged animals (Fig. 4D-E). Importantly dSetdb1 knockdown in OPNs, increased reporter activity in response to PQ in aged flies (Fig. 4D-E).

Single cell expression analysis specifically in cholinergic OPNs using Scope revealed a decrease in the UPR^MT^-associated chaperones Hsp60 and Hsc70-5 in aged OPNs (Fig. 3F-H), with constant levels of dSetdb1 and lower levels of Kdm2 (Fig. 4I-J). Importantly, this data suggests that the observed increase in trimethylation levels within OPNs may be associated with a decline in demethylase activity as flies age. Underlaying significant implications for the regulation of UPR^MT^ and the overall epigenetic landscape in aged neurons.

Having demonstrated that dSetdb1 is essential for the increase in H3K9me3 in aged flies, preventing the activation of UPR^MT^ specifically in OPNs, we next evaluated olfactory function. Knockdown of dSetdb1 only in cholinergic OPNs improved olfactory function in aged flies compared to controls. We then assessed if this improvement in olfactory function was dependent of UPR^MT^. To this end, dve or crc were knockdown specifically in cholinergic OPNs in dSetdb1-deficient flies. Importantly, knockdown of dve or crc reversed the maintenance of olfactory function induced by dSetdb1 knockdown, mirroring the olfactory capacity of control aged flies (Fig. 4L). Additionally, downregulation of dve and crc only in OPNs impaired olfactory function in young flies, to levels comparable to that of aged control flies.

### 5. Downregulation of dSetdb1 in OPNs restores age-associated mitochondrial morphological abnormalities and reduces mROS levels

As changes in UPR^MT^ activation can influence mitochondrial morphology and function, potentially triggering degenerative mechanisms, we investigated mitochondrial morphology in OPNs by expressing a mitochondrially targeted GFP. We examined mitochondrial morphology in three distinct compartments of OPNs, including cell bodies in the AL, the axonal tract, and the presynaptic terminal-enriched lateral horn (LH, Fig. 5A). In the AL of aged flies, a marked decrease in total mitochondrial volume was observed, along with increased mitochondrial fragmentation and sphericity. Notably, the targeted downregulation of dSetdb1 within OPNs mitigated these age-related changes, resembling the values observed in young control flies. Surprisingly, dSetdb1 knockdown also resulted in reduced mitochondrial fragmentation in young flies, suggesting a potential disruption of the mitochondrial network when compared with age-matched controls (Fig. 5B-F). Within the axonal tract, aged control axons exhibited a significant reduction in mitochondrial volume when compared to younger flies (Fig. 5G–H). Targeted knockdown of dSetdb1 effectively maintained mitochondrial volume. Lastly, no significant changes in mitochondrial parameters were observed in the LH of aged flies for both genotypes (Fig. 5I-J, and Supplementary Fig. 1).

**Figure 5.**
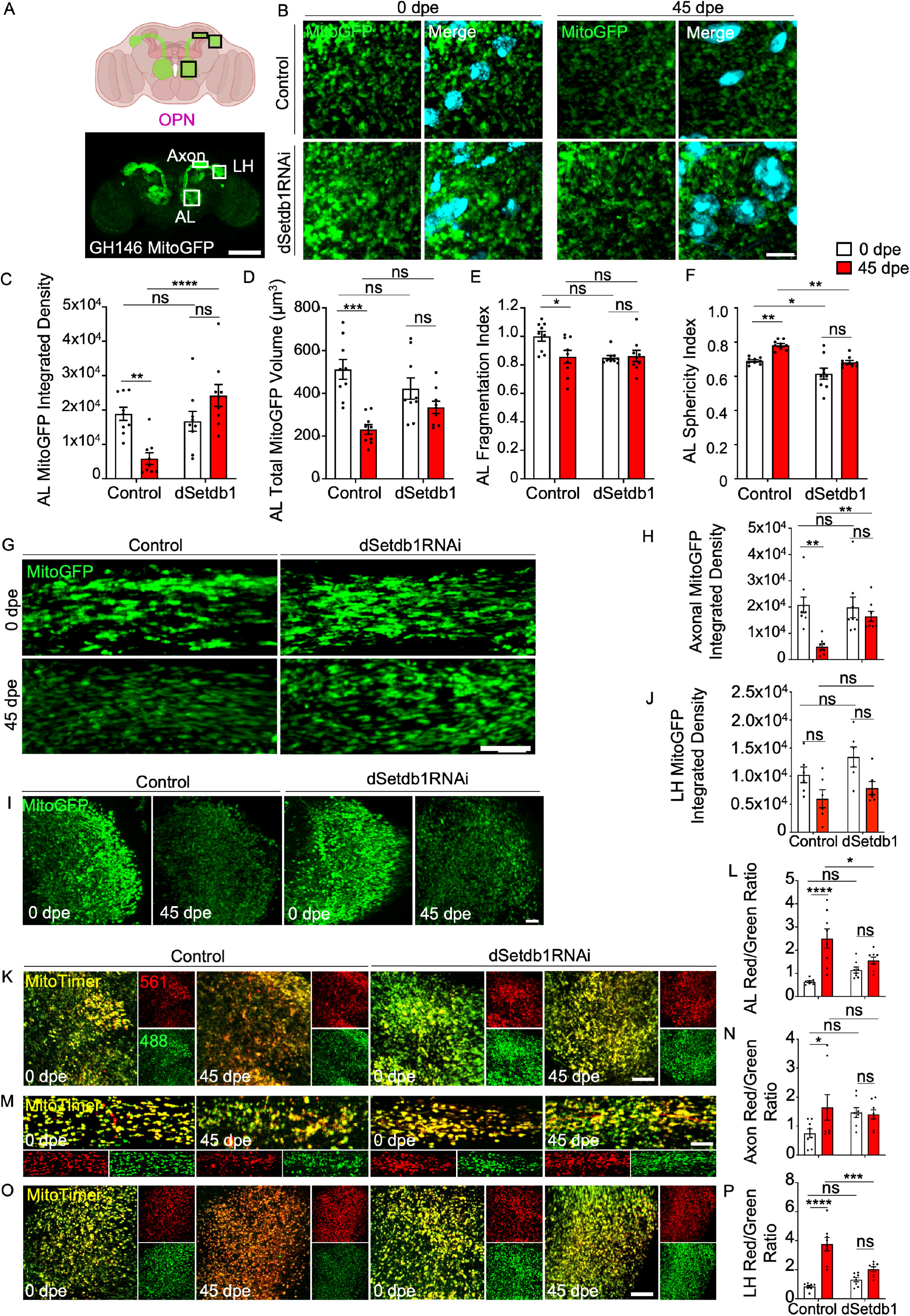
dSetdb1 Downregulation Mitigates Age-Related Mitochondrial Oxidation and Preserves Mitochondrial Morphology in OPNs. (A) The upper panel depicts the GH146-driven GFP expression in olfactory projection neurons (OPNs). The lower panel shows mitochondrial GFP reporter expressed in OPNs by the GH146 driver. Neurons possess their soma in the antennal lobe (AL) and project their axons to the synapsis enriched zone, the lateral horn (LH). Scale bar, 100 µM (B) Mitochondria labeled with GFP in the AL of OPNs of 0 and 45 dpe flies bearing the dSetdb1. Mitochondria in Green and nuclei in ToPro3 Scale bar, 5 µM. (C) Mitochondrial integrated density in the AL of 0 and 45 dpe flies with dSetdb1 knockdown analyzed via Two-way ANOVA with Bonferroni’s multiple comparisons test. Results: control flies showed a significant difference at 0 vs 45 dpe (p=0.0018, n=9), while dSetdb1 RNAi flies showed no significant change (p=0.0855, n=9). Comparison between Control vs dSetdb1 RNAi flies of 0dpe (p>0.9999, n=9), and 45 dpe (p<0.0001, n=9). (D) Analysis of total mitochondrial volume in the AL of OPNs with dSetdb1 knockdown compared to controls at 0 and 45 dpe. Two-way ANOVA with Bonferroni’s multiple comparisons test indicates a decrease in mitochondrial volume in control flies from 0 to 45 dpe (p<0.0001, n=9). The dSetdb1 RNAi of 0 vs 45 dpe flies did not show a change in volume (p=0.2387, n=9). (E) AL mitochondrial fragmentation index of images shown in B. Two-way ANOVA Bonferroni’s multiple comparisons test for mitochondrial fragmentation index in control flies from 0 to 45 dpe (p=0.0145, n=9) and dSetdb1 RNAi flies showed no significant change (p>0.9999, n=9). Control vs. dSetdb1 RNAi at 0 dpe (p=0.0101, n=9); no significant change at 45 dpe (p>0.9999, n=9). (F) AL sphericity index of mitochondria from images shown in B. Graph shows results of Two-way ANOVA with Bonferroni’s multiple comparisons test. For 0 vs 45 dpe, control flies (p=0.0019, n=9) and dSetdb1 RNAi flies (p=0.0257, n=9). At 0 dpe, control vs dSetdb1 RNAi flies (p=0.012, n=9), and at 45 dpe (p=0.0008, n=9). (G) Representative images of Axonal mitochondria in the green of 0 and 45 dpe flies bearing the knockdown of dSetdb1. Scale bar, 5 µm. (H) Axonal MitoGFP integrated density; control increase (p=0.0005, n=9), dSetdb1 RNAi (p=0.754, n=9). (I) LH MitoGFP confocal images. Scale bar, 10 µm. (J) LH MitoGFP integrated density; control (p=0.1197, n=9), dSetdb1 RNAi (p=0.0751, n=9). (K, M, and N) Representative images of 0 and 45 dpe *Drosophila*’s AL, Axonal, and LH showing GH146 GAL4;UAS-MitoTimer, an in vivo mitochondrial oxidation reporter. Unoxidized roGFP-labeled mitochondria appears in Green (488) and oxidized mitochondria appears in red (561) and Merge (Yellow). Scale bar, 5µm. (L) AL Red/Green integrated density ratio), the 0 vs 45 comparison reports a significant oxidation increase in control flies (p<0.0001, n=8), while dSetdb1 RNAi flies show a non-significant change (p=0.4359, n=8). (N) Axonal Red/Green integrated density ratio, 0 control flies shows an increase in oxidation at 45dpe (p=0.0447, n=8), with no changes in dSetdb1 RNAi flies (p>0.9999, n=8). (P) LH Red/Green integrated density ratio shows a oxidation increase in 45 dpe control flies (p<0.0001, n=8), while dSetdb1 RNAi flies do not display change (p=0.1263, n=8). White and red bars represent 0 and 45 dpe flies, respectively. n= Independent AL of *Drosophila* brain. P-value: **** p < 0.0001; *** p < 0.001; ** p < 0.01, * p<0.05 and ns > 0.05.

We next assessed mitochondrial oxidation levels as a surrogate marker of mitochondrial function^30,31^. To this end, we employed the UAS-MitoTimer construct, a mitochondrial oxidation reporter^31,32^. This tool relies on the expression of a green fluorescent protein that transitions to red fluorescence upon oxidation^31^. Compared to young control flies, we observed an increase in mitochondrial oxidation in older control flies within the AL (Fig. 5K-L), axonal tract (Fig. 5M-N), and the LH (Fig. 5O-P). Remarkably, downregulation of dSetdb1 in OPNs reversed the age-related mitochondrial oxidation in the three analyzed neuronal regions. Our results indicate that the downregulation of dSetdb1 in OPNs, activating UPR^MT^ during aging, not only reverses age-associated changes in mitochondrial morphology but also effectively prevents the accumulation of mitochondrial oxidation.

### 6. Epigenetic regulation of UPR^MT^ by dSetdb1 modulates age-dependent neurodegeneration of OPNs

Loss of neuronal function is often linked to degenerative structural changes in the neuronal circuit^33,34^, a phenotype extensively associated with mitochondrial dysfunction. Therefore, we investigated whether UPR^MT^ activation regulates neuronal integrity throughout aging in OPNs. To assess the impact of UPR^MT^ modulation in neuronal integrity, we first assessed changes in neuronal number throughout aging by counting nuclei from GFP-positive OPNs. In control flies, aging resulted in a significant decrease in the number of OPNs. Remarkably, this age-related neuronal loss was prevented in flies where dSetdb1 was downregulated only in OPNs (Fig. 6A-B). We next evaluated axonal integrity GFP-labelled OPNs. In aged control flies, a reduction in axonal integrated density was observed when compared to young control flies, consistent with the previously noted decrease in the total number of neurons. Notably, dSetdb1 knockdown protected against the decline in axonal integrity of OPNs associated with aging (Fig. 6C-D).

**Figure 6.**
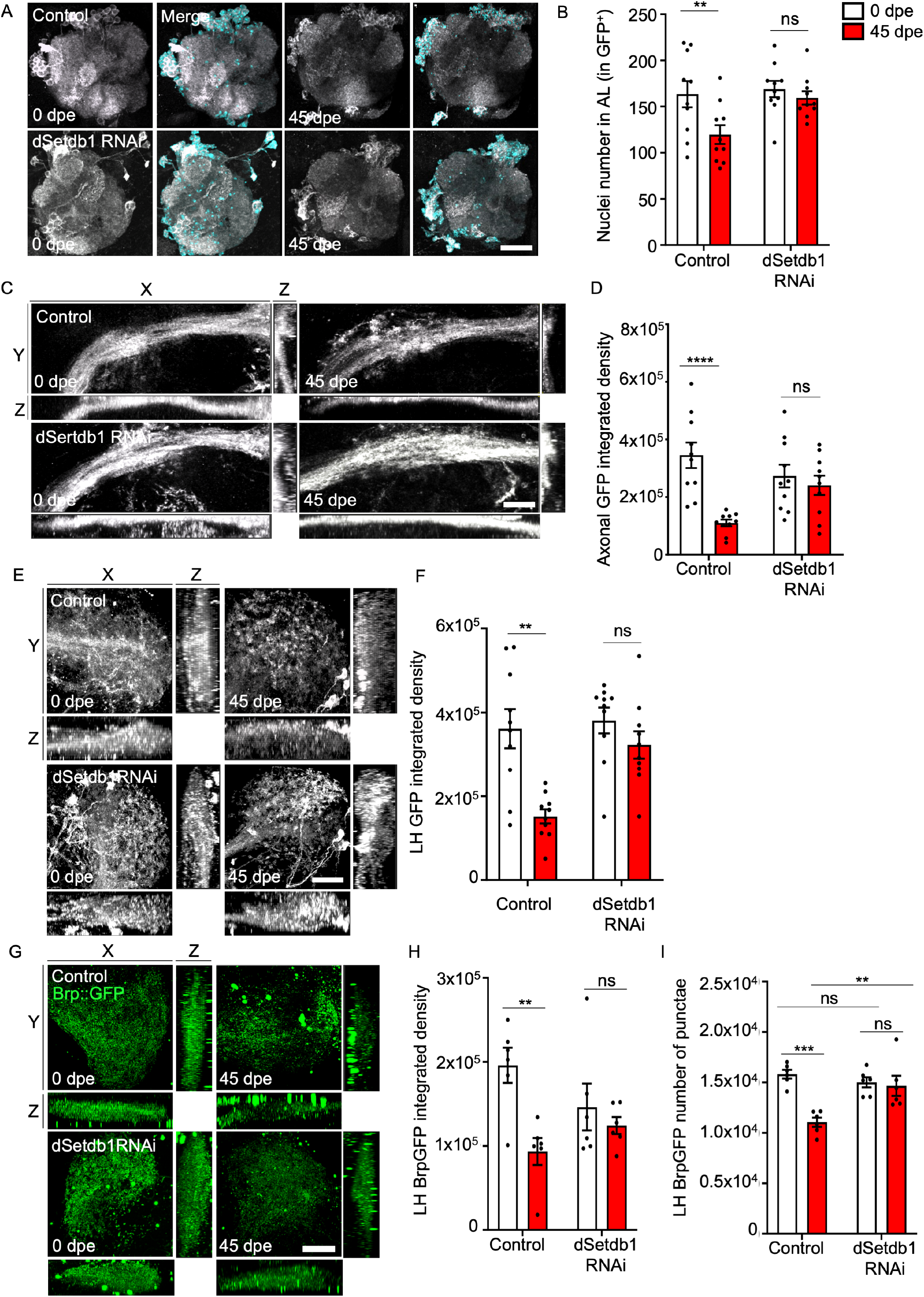
dSetdb1 Knockdown Preserves Neuronal Integrity and Synaptic Density in the Aging *Drosophila* OPNs. (A) GFP-labeled olfactory projection neurons (OPNs) in the AL of 0 and 45 dpe control flies and flies with knockdown of dSetdb1 and dve. GFP-positive OPNs are gray, and the right panel shows merged channels with nuclei labeled with ToPro3 in cyan. Scale bar, 20µm. (B) Quantification of nucleus count in GH146-positive OPNs. The graph shows Two-way ANOVA with Bonferroni’s multiple comparisons between 0 and 45 dpe shows that control flies had a significant decrease in nucleus count (p=0.0233, n=10). dSetdb1 RNAi (p>0.9999, n=10) Control vs. dSetdb1 RNAi of 45 dpe, (p=0.0398, n=10). (C) Orthogonal view of the 3D reconstruction of the distal axonal tract of OPNs tagged with GFP. Panel shows show the combination of axis Y and X, Z and X, and Z and Y. Panel shows axons from control flies, knockdown flies for dSetdb1. Scale bar, 5µm. (D) Quantification of axonal integrated density of axons shown in C. Two-way anova multiple comparison between Control flies show a significant decrease in axonal integrated density from 0 to 45 dpe (p<0.0001, n=10). dSetdb1 RNAi (p>0.9999, n=10). (E) Representative images of orthogonal view of the 3D reconstruction of GFP-tagged OPNs in the LH. Images show the combination of axis Y and X, Z and X, and Z and Y. Panel shows LH from 0 and 45 dpe control flies, knockdown flies for dSetdb1 and dve. Scale bar, 10µm. (F) Quantification of GFP integrated density in the LH of images shown in E. Two-way ANOVA with Bonferroni’s multiple comparisons test shows a significant decrease in GFP volume in the LH of control flies from 0 to 45 dpe (p<0.0001, n=10). And no change for dSetdb1 RNAi (p>0.9999, n=10). (G) Orthogonal view of representative 3D reconstruction images of Brp::GFP-labeled presynaptic densities in LH of 0 and 45 dpe flies bearing the dSetdb1 GH146 knockdown. Scale bar, 20µm (H Quantification of BrpGFP integrated density of images shown in G. LH Brp::GFP integrated density, there was a significant decrease in Brp::GFP integrated density in control flies from 0 to 45 dpe (p=0.0031, n=6), while dSetdb1 RNAi flies did not show a significant change (p=0.8823, n=6). (I) Quantification of number of presynaptic densities labeled with BrpGFP in the LH of flies bearing the dSetdb1 knockdown shown in G. Control flies showed a reduction in the number of presynaptic densities labeled with Brp::GFP from 0 to 45 dpe (p<0.0001, n=6), and dSetdb1 RNAi no significant change (p>0.9999, n=6).White and red bars represent 0 and 45 dpe flies, respectively. n = independent fly brain. P-value: **** p < 0.0001; *** p < 0.001; ** p < 0.01, * p<0.05 and ns > 0.05.

It has been previously demonstrated that the age-associated decline in olfactory function in *Drosophila* is associated to the loss of synapses in OPNs^5^. Thus, we focused on the LH, a region enriched in presynaptic connections of OPN neurons^5,35^. Control flies exhibited a significant decrease in LH integrated density throughout aging, which was prevented by downregulation of Setdb1 in OPNs (Fig. 6E-F). This data suggests that H3K9-dependent UPR^MT^ activation plays a crucial role in maintaining neuronal integrity within this presynaptic enriched region. To gain a more detailed insight into synaptic zones, we employed the Brp::GFP reporter, a fusion protein that specifically accumulates in presynaptic buttons, facilitating visualization and quantification of presynaptic puncta^36^. Our analysis revealed that both total volume and number of GFP puncta in the LH of aged control flies were reduced compared to their younger counterparts (Fig. 6G-I). When dSetdb1 was downregulated in OPNs, we observed a decrease in this age-dependent integrated density decline. Notably, there was no significant difference between the number of Brp::GFP-positive puncta in young and aged dSetdb1 knockdown flies (Fig. 6I).

These findings collectively demonstrate that the targeted downregulation of dSetdb1 plays a causal role in preserving neuronal numbers and axonal integrity in OPNs. This intervention not only maintains presynaptic densities but also actively contributes to the demethylation of H3K9 and the activation of UPR^MT^, which are integral to the preservation of olfactory function during aging. By modulating these key processes, our results establish a direct link between the epigenetic regulation by dSetdb1 and the mitigation of age-related neurodegeneration in OPNs, underscoring the potential of targeted epigenetic interventions in maintaining neural health in olfactory system.

## Discussion

The UPR^MT^ plays a critical role in preserving mitochondrial homeostasis^37^. Therefore, any change in its activation potential, whether caused by physiological or pathological factors, could impact mitochondrial function^38–41^. Although it is widely accepted that olfactory function declines with age, diverse factors have been associated with this age-related impairment^42–46^. Our findings indicate that the age-dependent decline of olfactory function in *Drosophila* is associated to a decrease in the activation capacity of the UPR^MT^ in olfactory neurons. Crucially, the reduction in UPR^MT^ activation is linked to an increase in H3K9me3, which is dependent on the methylation activity of dSetdb1. Importantly, targeting this methylation process can effectively prevent age-related neuronal degeneration and restore the loss of olfactory function associated with aging.

Our study uncovers a novel aspect of epigenetic regulation in the aging process, emphasizing the specific role for dSetdb1 in modulating H3K9me3 levels and suppressing of the UPR^MT^. While previous research has established the impact of H3K9 methylation on the UPR^MT^, primarily through the actions of the demethylases JMJD 3.1 and JMJD 1.3, as well as methyltransferases such as MET-2 (dSetdb1 orthologue), SET6, BAZ2 in worms and mice respectively^14,15,41^, the specific function of Setdb1 in this context remained unclear. Our analysis indicates that in the AL of *Drosophila* brain, the absence of dSetdb1, also known as *eggless*, leads to a reduction in H3K9me3 in aged flies, suggesting its role as a tri-methyltransferase that acts as an epigenetic modulator of UPR^MT^ during aging. This finding aligns with evolutionary conservation in epigenetic regulation across species. Supporting this, research shows that in *C. elegans*, knocking out met-2 also reduces H3K9me3, suggesting a conserved mechanism^47^. However, previous research on the role of met-2 in UPR^MT^ regulation only focused on H3K9Me2 and did not extended their examination to H3K9me3^14^ Additionally, the role of SET6 in tri-methylating H3K9 and reducing UPR^MT^ in mice further highlights a possible conserved epigenetic pathway influencing aging across distinct species^41^.

Existing research underscores the beneficial role of UPR^MT^ in maintaining cellular integrity, primarily through maintaining mitochondrial function, a critical factor for healthy aging^48–50^. Indeed, the overexpression of the histone demethylases JMJD-1.2 and JMJD-3.1 extend lifespan in *C. elegans*^40^. Conversely, reduced expression of UPR^MT^ nuclear effectors ATFS-1, UBL-5, and DVE-1, as well as demethylases JMJD-1.2 and JMJD-3.1, compromises lifespan and cellular viability^15,48,51–53^. Recent work reveals the fine tunned regulation of UPR^MT^ along aging, particularly through the H3K9 methyltransferase SET-6 and the epigenetic reader BAZ-2, which modulate gene expression related to mitochondrial health and stress responses, essential for neuronal viability^54^. Importantly, beyond the impact on lifespan, UPR^MT^ regulation has profound implications for neuronal function. In *C. elegans*, mitochondrial function was found to influence pharyngeal pumping (eating) and defecation rates, crucial for lifespan^50^. Additionally, deficits in dopamine-dependent behaviors were observed in pdr-1 and pink-1 mutants, indicative of neuronal dysfunction without neuronal loss. This dysfunction is exacerbated by the downregulation of atfs-1, which is critical for UPR^MT^ ^52^. In a mammalian context, Baz2b ablation enhanced mitochondrial function in the hippocampus and cerebellum in older mice, suggesting its role in modulating age-related cognitive decline^41^. This was accompanied by improved performance in locomotion, reflecting preserved motor functions in aged mice. Beyond these, UPR^MT^ regulates hippocampal neural stem cell aging, with implications for cognitive functions^55^ and affects skeletal muscle aging, as exercise improves coordination of UPR^MT^ and mitophagy in aging skeletal^56^. Moreover, it plays a critical role in fertility and reproductive aging^57^ suggesting that UPR^MT^ disruption can pivot healthy aging towards pathological states. Our data indicate that reducing H3K9me3 levels during aging to enhance UPR^MT^ activation is beneficial for olfactory function and support the prevailing hypothesis that modulation of epigenetic regulators, which suppress UPR^MT^ transcriptional activation, constitutes a viable therapeutic approach for ameliorating mitochondrial dysfunction associated with aging.

Brain aging exhibit distinct regional variations across multiple levels, including gene expression, organelle, and neuronal function^58–60^. Our analysis of gene expression from single cell studies reveals neuronal-specific transcriptional expression for H3K9-regulating enzymes such as dSetdb1, utx, and kdm2 in aged vAChT, vGlut, and Gad1 neurons. It has been previously demonstrated that H3K9me3 levels are not uniform across the brain, varying based on brain regions or neuronal types^61–63^. Neurodegenerative conditions, characterized by the selective degeneration of specific neurons and their projections, exhibit differential neuronal vulnerability that is intricately linked to variations in neuronal morphology, activity patterns, and gene expression profiles within these affected structures^64–66^. Our data suggests that the age-dependent reduction in H3K9 demethylation enzymes within specific neuronal populations may contribute to differential neuronal vulnerability along aging, which deserves further exploration.

The increase we had shown in the levels of H3K9me3 in the brains of aged fruit flies aligns with prior studies reporting a rise in H3K9me3 in aged *Drosophila* heads^33^. Studies performed in *C. elegans* have revealed that as age progresses, there is an increase in the expression of the H3K9me3 methyltransferase SET-6 and the epigenetic reader BAZ-2. Remarkably, inhibiting their expression has been linked to preservation of pharyngeal pumping in these organisms^41^. In mice, administering an inhibitor for the histone methyltransferase SUV39H1, which is responsible for the trimethylation of H3K9, was found to mitigate age-associated cognitive decline and augment dendritic spines in the hippocampus^67^. Such findings support the notion that diminishing H3K9me3 levels might enhance functionality of different brain modules during aging. While our findings suggest that reducing hypermethylation in aging could potentially enhance UPR^MT^ response, it is critical to acknowledge that methylation processes also govern a vast of other cellular and neuronal functions^68^. For instance, histone methylation plays a crucial role in gene expression regulation, cellular differentiation, and even neuronal activity^69,70^, all of which could be inadvertently impacted by broad-spectrum epigenetic interventions. This pleiotropic nature of methylation underscores the importance of a targeted approach.

Mitochondria play a central role as primary sensors for degenerative stimuli^71^. With aging, neurodegeneration is often preceded by mitochondrial dysfunction, which manifests as morphological changes, including swelling, fragmentation, reduced volume, and increased oxidative stress^72–80^. Consistent with prior studies in aged flies^5^, we did not observe an increase in fragmentation within axons of the OPNs and lateral horn (LH), which could be attributed to the specific neuronal types under investigation^81^. Interestingly, our observations align with those from other studies^5^, as we identified an age-related deterioration in *Drosophila’s* olfactory circuits, which coincides with a rise in oxidative mitochondria. Current research underscores the central role of UPR^MT^ activation in orchestrating mitochondrial morphology and function^82–86^. Importantly, our study demonstrated that genetic inhibition of dSetdb1 restored youthful levels of H3K9me3, enabling UPR^MT^ activation to restore mitochondrial morphology and oxidative status. This maintenance of cellular viability through UPR^MT^ activation parallels findings from a recent study, which revealed that mild mitochondrial dysfunction-dependent UPR^MT^ activation, protects cardiomyocytes against cardiac ischemia-reperfusion injury in a mouse model^9895^. Also, NAD+ activation of the UPR^MT^ rejuvenated muscle stem cells in aged mice^82^, highlighting the role of UPR^MT^ in altering aging markers in stem cells and extending lifespan.

Our research highlights the crucial role of UPR^MT^ regulation in the age-related decline of olfactory function. Focusing in the olfactory system in aged *Drosophila*, we have demonstrated detrimental effects of epigenetic changes on mitochondrial function, impacting neuronal survival. The importance of olfaction extends beyond sensory perception, with human olfactory processing is intricately linked to emotions and memories, mediated by the limbic system and cerebral cortex^87,88^. A compromised sense of smell is not only associated with depression in a significant number of cases^89^ but also frequently precedes the onset of age-related neurodegenerative diseases such as Alzheimer’s^90,91^ and Parkinson’s disease^92,93^. Our findings underscore the need for further exploration of the UPR^MT^ pathway and its epigenetic regulation as a potential target for developing interventions to mitigate the decline in neuronal function associated with the aging process.

## Materials and Methods

### *Drosophila* Strains and Culture

Strains carrying the following transgenes were obtained from the Bloomington *Drosophila* Stock Center (BDSC) from Indiana University: UAS-Mito-GFP (BL#8443, encodes the 31 amino acid mitochondrial import sequence from human cytochrome C oxidase subunit VIII fused to the N-terminus of the Green fluorescent protein), UAS-MitoTimer (BL#57323, encodes a mitochondrial targeting sequence with a roGFP which turns its fluorescence from green to red when oxidized), Elav-Gal4 -Gal4 (BL#485), Actin-Gal4 (BL#9431), GH146-Gal4 (BL#1104), GH146,GFP (BL#36500); RNAi lines for UPR^MT^ genes including crc (BL# 25985), dve (BL#26225), ubl (BL#65893), and RNAi lines for UPR^MT^ associated H3K9 methylation enzymes dSetdb1 (BL# 31352), dSetdb1 loss of function (BL#30566), Utx (BL#34076), and Kdm2 (BL#33699). Wild-type flies used as controls are Canton S (BL#64349), and RNAi Control flies from the Transgenic RNA Interference Project (TRIP) (BL#35787). Only female flies were used for all experiments to avoid genetic variation and aggressive behavior from males. All fly stocks were maintained on a standard *Drosophila* medium which consisted of 112.5g of Molasses, 35g of dry yeast, 90g of corn flour, 9g of agar, 2.5g of Tegosept diluted in 10ml of ethanol 95%, and 6ml of propionic acid per 1L of water; at 25°C and under a circadian cycle of 12h of light and 12h of darkness.

### Treatment with Mitochondrial Stressor Paraquat

Experiments requiring mitochondrial stressor paraquat (Sigma, 36541) were performed by supplementing standard *Drosophila* medium with 100µl of paraquat diluted in dH2O at 10µM. After PQ addition, vials must be airdried before use. Groups of flies were exposed to paraquat-supplemented medium for 48hrs before experiments were conducted.

### Olfactory functional assay

Olfactory T-maze was used to perform the olfactory behavioral test based on Hussain et al. 2018. Briefly, 15 flies are presented with an abrasive odor 0.1M of Hydroxychloroquine, 3-octanol (Sigma, W358118), or a pleasant odor of 2.3-butanedione (Sigma, B85307) at the end of one arm of the T-maze and at the end of the opposite arm flies are exposed to control solution (vehicle only). Flies have 60 seconds to discriminate between odors and go to an arm of the T-maze. At the end of the 60 seconds, an image is acquired, and flies in both arms are counted. The olfactory preference index consists of ((Flies in Experimental Odor – Flies in Vehicle Odor) / (Total flies in the experiment)). The olfactory preference index is calculated for every trial, and every n in the graph corresponds to the mean of 5 trials of 15 flies each. For statistical analysis when comparing genotype and treatment, two-way ANOVA tests were performed with analysis of variance and Dunnet multiple comparisons for more than two groups using GraphPad Prism 6. P-value: **** p < 0.0001; *** p < 0.001; ** p < 0.01, * p<0.05 and ns > 0.05.

### Protein Quantification and western blotting

For immunostaining of H3K9me3, samples were prepared as described previously3. Briefly, samples were frozen in liquid nitrogen and ground to a fine powder using a pestle fitted to a 1.5ml Eppendorf Centrifuge tube filled with RIPA buffer, which included 50nM Tris, 150 mM NaCl, 1 mM EDTA, 0.1% of Nonodet P-40 (NP-40), 0.25% of Sodium Deoxycholate, and 0.02% of sodium azide in ddH20. RIPA buffer was supplemented with phenylmethylsulphonyl fluoride (PMSF – Sigma) and a protease inhibitor cocktail (Sigma, P8340). Homogenized samples were incubated at 4°C for 1h and sonicated by sonicator (QSonica) at 40% of equipment maximal amplitude with 3 pulses in 1 min. Sonicated Samples were centrifuged at 500g to pellet the debris, and all supernatant was transferred to a new tube and then centrifuged for 14 min at 13000g at 4C. The upper soluble phase was transferred to a new 1.5ml Eppendorf for membrane and plasma proteins and pellet, and 200ul of liquid phase was kept for nuclear proteins. Pellet was dissolved in the liquid phase by pipetting. Quantification of samples was performed using the Pierce BCA Protein Assay Kit (Thermo Scientific, 23225) under the manufacturer’s instructions. Samples were boiled in SDS sample buffer for 15 min, separated on an SDS-PAGE gel, transferred, and revealed using BioRad TransBlot and ChemiDoc, respectively. Primary antibodies used were rabbit anti-H3K9me3 1:2000 (Abcam, ab8898), and loading control anti-Tubulin 1:1000 (Thermofisher, MA1-744). Secondary antibodies were anti-rabbit conjugated with Horse Radish Peroxidase (HRP) 1:1000 (Thermofisher). The stained membranes were briefly incubated in luminol and scanned using ChemiDoc (BioRad). One biological replicate corresponded to a homogenized solution of 20 fly heads minimum. The normalized H3K9me3 levels were calculated by normalizing the ratio of H3K9me3 and loading control to that of young control samples. The significance of the interaction between genotypes and time was calculated by a two-way ANOVA test with Dunnet multiple comparisons using GraphPad Prism 9. P-value: **** p < 0.0001; *** p < 0.001; ** p < 0.01, * p<0.05 and ns > 0.05.

### Dissection of adult *Drosophila* brains and confocal microscopy

Flies of the desired genotype were collected in groups of 15 and placed in vials containing *Drosophila* medium. Flies were anesthetized using a CO_2_ pad. Using Dumont forceps #5, flies were held from the thorax and dipped in the sylgard petri dish filled with cold PBS. Brains were isolated by removing the exoskeleton from the flýs head and carefully removing the esophagus and air sacs of the flies’ brains as previously described^4^. *For live imaging of endogenous fluorescence signal* (Fig. 1A, C, E, 2A, 3C, E, 4H, 6A, C, E and 8C) brains were fixed in 2% paraformaldehyde with 0.1% Triton X-100 (Sigma, T9284). for 20 minutes and then changed to a 4% paraformaldehyde, 0.1% Triton X-100 solution for 20 more minutes, then washed for 10 min 3 times in PBS Triton X-100 at 0.1% followed by 3 quick washes with PBS only. Brains were mounted in VectaShield antifade mounting medium (Vector, H1000) for later visualization in an SP8 confocal microscope. For all experiments, all brains were imaged on the same day. *For immunostaining* (Fig. 4J, 5A,B,G,I, 7A,C, and 8A), brains were dissected as described previously5. Brains were fixed for 1h at room temperature in 4% PFA with 0.5% Triton X-100, washed 3 times for 10 minutes with PBS with 0.5% Triton X-100, and blocked for 1h with Normal goat Serum (NGS) (Cell Signaling, 5425S) at 5% in PBS, 0.5% Triton X-100 and stained overnight at 4°C with primary and after 3 washes in PBS, 0.5% Triton X-100 with secondary antibodies using the same conditions. The secondary antibody was washed 6 times for 10 min each with PBS, 0.5% Triton X-100 a 3 quick washes in PBS before mounting. Washed brains were placed in a stripe of 15 to 20 µl of Vectashield antifade mounting medium (Vector, H1000) on cover glass (Deltalab, D10004), and imaging was performed in confocal microscope SP8 using the 63x objective with digital zoom necessary for desired resolution. Fluorescence intensity for each channel was adjusted using control flies to the point that no saturation was observed, then the same parameters were used for all images. Images of antennal lobe sections of *Drosophila* brains and OPNs were taken at a depth of 10µm using a Z stack separation of 0.6µm. Primary antibodies used were anti-GFP (Invitrogen, 1:1000), rabbit anti-H3K9me3 (Abcam, ab8898; 1:500), and secondary antibody donkey anti-Rabbit 555 (Thermofisher, 1:1000).

### Image quantification

For image quantification 3D reconstruction of labeled structures was performed in Imaris Software. The surface of the desired signal was rendered. And the following parameters where quantified. *Integrated density:* This parameter represents a cumulative metric of the fluorescence signal within a specified region, denoting the aggregate of signal intensity and its spatial distribution. It is computed by multiplying the average fluorescence intensity by the volume of the signal-bearing domain, thereby yielding a singular value that encapsulates both the concentration and extent of the fluorescent activity. This approach ensures normalization for variations in the volume of the assessed Region of Interest, thereby facilitating a more accurate comparative analysis across samples.. *Mitochondrial Morphology:* Mitochondrial changes during aging in OPNs were analyzed by rendering the surface of mitochondrial reporter mitoGFP expressed specifically in the OPNs. Total mitoGFP volume (µm^3^), mitoGFP puncta number, mitoGFP fragmentation index which corresponds to volume (µm^3^) per area, (µm^2^) mitoGFP sphericity index, mitoGFP average size (µm^3^), and integrated density were analyzed. *Mitochondrial Oxidation:* Mitochondrial oxidation was assessed using MitoTimer, a fluorescent protein that shifts from green to red upon oxidation. The analysis involved processing both the green and red channels and determining their ratio. In this context, oxidized mitochondria appeared red, while healthy mitochondria were green, following the methodology outlined in a previous study^31,32^. *Nuclei number and H3K9me3 quantification:* For quantification of the specific signal in Olfactory Projection Neurons, a surface of GFP-labeled OPNs was rendered, then To-Pro3 labeled nuclei signal was masked and the surface rendered, and finally, the signal for H3K9me3 in the nuclei of OPNs was quantified as described previously^5^. *Neuronal Integrity:* Neuronal degeneration of OPNs was analyzed by rendering the surface of GFP-labeled OPNs in AL, the distal part of the axon, and axonal terminals in the LH, Integrated density was calculated to determine the amount of signal intensity. *Presynaptic puncta Quantification:* For quantifying presynaptic puncta, we employed the Brp::GFP reporter, a fusion of Brunchpilot and GFP. This reporter accumulates in presynaptic buttons, allowing visualization. After rendering the GFP surface, we applied a mask to the GFP channel and counted the puncta, using a threshold ratio of 300µm in the masked channel to ensure accuracy as described previously^36^. Data was plotted and analyzed using GraphPad Prisms 9 Software. To compare the interaction between age and genotype/treatment, two-way ANOVA and for more than two groups, analysis of variance with Dunnet multiple comparisons was performed using GraphPad Prism 9. P-value: **** p < 0.0001; *** p < 0.001; ** p < 0.01, * p<0.05 and ns > 0.05.

### Single Cell RNA-seq Data analysis

To analyze gene expression in different cells population between the *Drosophila* aging brain, we used SCope (http://scope.aertslab.org) or the “ScopeLoomR” package in R with the scRNA-seq data “Aerts_Fly_AdultBrain_Filtered_57k.loom” under accession code GEO:GSE107451. We compared the AUC (Area Under the Curve) values derived from SCope, which indicate the activity levels of genes under regulons across diverse cellular populations. These values, reflects the combined activity of gene sets regulated by specific transcription factors, allowing to infer changes in gene expression. By assessing the AUC values for specific genes of interest—namely Hsp60A, Hsc70-5, dSetdb1, Utx, and Kdm2—we could quantitatively evaluate their activity in their respective regulon within distinct neuronal populations, including cholinergic (vAChT), glutamatergic (vGlut), GABAergic (Gad1), and olfactory projection neurons (OPN). This approach allowed us to quantify a proxy for gene abundance and enabling a nuanced understanding of the regulatory mechanisms at play. For a comprehensive understanding of the technical underpinnings and applications of SCope in single-cell transcriptomics, we refer to the work by Davie et al. (2018)^28^, which established a single-cell transcriptome atlas of the aging *Drosophila* brain. All data was tabulated in R and then plotted using GraphPad Prism 9 for statistical analysis.

### RNA extraction, RT-PCR and qPCR

Total RNA was extracted either from 2 whole flies or a minimum of 20 heads using a typical trizol extraction following the next steps: tissue was placed on a 1.5 ml Eppendorf tube and 300 µl of trizol reagent were added before homogenization with a plastic pestle 60 times. At this point sample can be stored at −80°C. 700 µl of trizol were added and samples were centrifuged at 13000 rpm for 10 min at 4°C to pellet the tissue debris. Supernatant was transferred to a fresh 1.5 tube. 300ul of chloroform was added, and samples were shaked vigorously by hand 15 times. Then incubation of the samples at room temperature for 5 min. Samples were centrifuged at 13,000 rpm for 15 min at 4°C. The upper phase was carefully transferred to a new RNAse-free tube, without touching the interphase. 700uL of isopropanol were added to precipitate the RNA and samples were incubated for 5 minutes at room temperature or 1 hour at −20°C. Centrifuge at 12,000 rpm for 15 min at 4°C. Discard the supernatant and RNA pellet was washed with 1 ml of ethanol 70% prepared with miliQ quality H2O. Centrifuge at 13000 rpm for 10 min at 4°C. Air dry the pellet briefly, resuspend the pellet in an appropriate volume of MiliQ water (20 to 50 µl). The RNA concentration was measured for each sample in duplicate using a NanoQuantMultiskan spectrophotometer and the purity of the sample was evaluated using the 260/280nm absorbance ratio. Reverse transcriptase-PCRs (RT-PCR) were carried out using the iScript RT Kit for cDNA synthesis (BioRad, 1708841). Where the final mix include 1000ng of RNA sample and 5µL of iScript RT-PCR mix with reverse transcriptase, final mix was carried to a final volume of 20ul. The cDNAs obtained were then used as a template for real-time PCRs carried out using the Eva Green qPCR Master Mix (SolisBioDyne, 08242510.6) in a StepOne Plus Machine (Applied Biosystem). The final PCR mixture (20ul) contained 1ul of cDNA, 4 ul of 5xFirePol MasterMix, 0.2umoles of each primer and was carried out to the final volume of 20ul. The thermal profile used for the reaction included a 2-minute heat activation of the enzyme at 95°C, followed by 35 cycles of denaturation at 95°C for 15 seconds and annealing/extension at 58°C for 60 seconds, followed by melt analysis ramping at 58-90°C. Negative control was conformed of water instead of cDNA and were included in each plate. Relative transcript levels were assessed using the Comparative CT method and expression values were normalized to 28S ribosomal expression, used as an internal control. One biological replicate corresponded to a homogenized solution of 20 fly heads minimum. For statistical analysis when comparing genotype and treatment, two-way ANOVA tests were performed with analysis of variance and Dunnet multiple comparisons for more than two groups using GraphPad Prism 6. P-value: **** p < 0.0001; *** p < 0.001; ** p < 0.01, * p<0.05 and ns > 0.05.

## Acknowledgments and Funding

This work was supported by grants from the Geroscience Center for Brain Health and Metabolism, FONDAP-15150012, Fondo Nacional de Desarrollo Científico y Tecnológico (FONDECYT) N° 1150766 (to FAC), Agencia Nacional de Investigación y Desarrollo (ANID) Subvención a la Instalación en la Academia PAI77180059 and FONDECYT Iniciación N°11200981 (to MS). We thanks Dr. Ramón Ramirez for his useful support on image processing with the Imaris software at the Center for Integrative Biology, Universidad Mayor de Chile.

## Supplementary Figure legends

**Supplementary Figure 1. Analysis of Mitochondrial Morphology in OPNs Unaffected by dSetdb1 Downregulation with Aging in Drosophila.** (A) Quantification of mitochondrial count in the antennal lobe (AL). (B) Average size of mitochondria in the AL. (C) Total mitochondrial volume in the AL. (D) Mitochondrial count in axonal regions. (E) Fragmentation index of axonal mitochondria. (F) Sphericity index of mitochondria in the AL. (G) Average size of mitochondria in axonal regions. (H) Mitochondrial volume in axonal regions. (I) Mitochondrial count in the lateral horn (LH). (J) Fragmentation index of LH mitochondria. (K) Sphericity index of LH mitochondria. (L) Average size of mitochondria in the LH. In panels A, D, and I, mitochondrial count remains unchanged by dSetdb1 downregulation across the age spectrum. In panels B and G, average mitochondrial size decreases with age in both control and dSetdb1 RNAi flies. In panel C, total mitochondrial volume is reduced in control flies with age but not significantly affected by dSetdb1 knockdown. In panels E, F, J, and K, the fragmentation and sphericity indices in axonal and LH regions show no differences between control and dSetdb1 RNAi flies, indicating that dSetdb1 does not affect mitochondrial morphology in these regions. Similarly, mitochondrial volume in axonal regions (H) and average mitochondrial size in the LH (L) are not affected by dSetdb1 knockdown. Data are presented as mean ± SEM; n=6, each n represents the number of independent biological replicates analyzed per group. Scale bars: 5 µm for axonal regions, 10 µm for AL and LH images. P-value significance: ns = not significant, * p < 0.05, ** p < 0.01, *** p < 0.001. Statistical significance was determined using two-way ANOVA with Bonferroni’s multiple comparisons test.

**Supplementary Figure 2. Schematic Representation of Age-Related Changes in H3K9 Methylation and Its Impact on UPR^MT^ and Neuronal Integrity in *Drosophila*.** In young organisms, mitochondria are challenged by various insults that lead to the accumulation of mitochondrial dysfunction, causing damage and activating a retrograde response from mitochondria to the nucleus. Ubl, crc and DVE are translocated to the nucleus, and DVE maintains an open chromatin state, allowing the binding of transcriptional modulators of the UPR^MT^. This event activates the transcription of chaperones and proteases to recover mitochondrial homeostasis and oxidation. During aging, trimethylation of H3K9 is a mark of heterochromatin associated with the repression of transcription. This trimethylated state of H3K9 does not allow the binding of the transcriptional modulators of UPR^MT^, dve, crc and ubl inhibiting the mitochondrial response to aging-causing damage. Thus, mitochondrial function persists and builds up in time-dependent manner, increasing mitochondrial oxidation and contributes to aging phenotypes, such as neurodegeneration marked by the reduction of OPNs, axonal volume, and presynaptic connections. Figure created with BioRender.com.

